# Camouflage functions of juvenile giant panda colouration in forest environments

**DOI:** 10.64898/2026.10.12.758461

**Authors:** Zixiang Li, Laura A. Kelley, Martin Stevens

## Abstract

The contrasting black-and-white coloration of the giant panda (*Ailuropoda melanoleuca*) has long attracted attention. Recent works suggest this contrasting pattern may generate camouflage-relevant visual effects. However, most previous studies have focused on adult pandas, whereas juveniles may face greater predation risk and may therefore represent a more ecologically relevant life stage for testing concealment hypotheses. We investigated whether juvenile panda coloration provides the potential for camouflage through background matching or disruptive coloration, and how this potential is shaped by vegetation cover, illumination and background position. We used life-sized juvenile panda models with natural and experimentally modified coloration by adjusting the relative proportions and spatial arrangement of white, black, and grey patches. Models were photographed in temperate woodland environments at ground level and against canopy backgrounds, under sunny and cloudy conditions, with and without vegetation occlusion. We quantified colour and luminance contrast, as well as edge-disruption metrics, using dichromatic canid and felid visual models. Natural juvenile panda coloration influences target-background contrast and edge-disruption, although these effects varies with vegetation, lighting, background height and visual model. Vegetation cover significantly reduced contrast and increased edge disruption metrics, while bright light condition only increased the metrics of edge disruption. Background position affected camouflage-relevant metrics in a context-and observer-dependent manner: under vegetation occlusion, canopy placement reduced chromatic contrast under felid-like vision, whereas ground-level placement reduced luminance contrast under both visual models. Alterations to patch extent and boundary arrangement modified camouflage-relevant metrics under specific environmental conditions, suggesting that the natural arrangement of pelage boundaries may enhance concealment under specific conditions. Overall, our findings support the hypothesis that juvenile giant panda coloration can produce visual effects consistent with potential camouflage, while highlighting the importance of habitat structure and viewing context in shaping camouflage.

## Introduction

The diversity and functionality of animal coloration have long been of interest as classical examples of evolution and natural selection (Darwin, 1872, Wallace, 1889). Camouflage is a widespread and common anti-predator strategy among animals, which rely on coloration to avoid detection or recognition by predators (Stevens and Merilaita, 2009, Cuthill, 2019). This anti-predator strategy includes many forms, such as background matching by blending body pattern and colours with the background (Stevens and Merilaita, 2009), disruptive coloration which hinders detection by breaking up the body shape or creating false outlines (Cuthill et al., 2005), masquerade involving resembling inanimate inedible objects (Skelhorn et al., 2010a), and using countershading to reduce self-shadows or 3D structure cues (Ruxton et al., 2004). While many camouflaged species exhibit low-contrast, background-matching coloration, some animals use bold high-contrast patterns, such as black-and-white stripes or patches, that appear conspicuous to human observers.

The giant panda (*Ailuropoda melanoleuca*) famously has high-contrast black-and-white pelage that has also long puzzled naturalists. Giant pandas live mainly in temperate high elevation forests in southwest China, where habitats are often characterised by strong variation in illumination, canopy shadows, dense vegetation and, in some areas or seasons, snow cover (Schaller, 1985, Loucks et al., 2003). Adult pandas typically exceed 50 kg after approximately 2.5 years of age, face limited predation risk (Schaller, 1985, Hu, 2001). By contrast, juvenile pandas are more vulnerable, and may climb trees to evade predators (Hu, 2001). Documented or potential predators include yellow-throated marten (*Martes flavigula*) (Harris, 2014), leopards (*Panthera pardus*), dholes (*Cuon alpinus*), brown bears (*Ursus arctos*), Asian black bears (*Ursus thibetanus*), and historically, wolves (*Canis lupus*) and tigers (*Panthera tigris*) (Morris and Morris, 1981, Schaller, 1985, Hu, 2001). Several of these predators are capable of climbing or attacking juveniles in structurally complex forest habitats, meaning that tree use may reduce but not eliminate predation risk. Moreover, because bamboo is nutritionally poor, pandas must allocate more time to foraging, which may reduce the parental care time and increasing the predation risk to offspring (Schaller, 1985). Juvenile pandas may therefore benefit from multiple anti-predator strategies, including behavioural avoidance and camouflage.

Early explanations for panda colouration proposed that white fur helps conceal pandas against snowy backgrounds (Morris and Morris, 1981, Schaller, 1985), or dark patches intersecting the body outline could generate disruptive effects (Caro, 2009). More recently, Caro et al. (2017) used a comparative phylogenetic approach and compared the pelage regions and their putative function across carnivores, found that white body regions were associated with snowy environments, whereas black regions were consistent with concealment in shaded areas. Nokelainen et al. (2021) further analysed adult panda photographs from camera traps, reported that black fur may blend with tree trunks, white fur with snow and foliage, and intermediate off-white pelage tones matched rocks and soil. They also found high estimates of potential edge disruption, particularly at viewing distances above approximately 50 m for dichromatic observers. However, the study was based mainly on adult pandas which face relative low predation risk, and long viewing distances may be less directly applicable to cluttered forest habitats where vegetation often restrict visibility (Lang et al., 2021a, Lang et al., 2021b). It therefore remains unclear whether juvenile pandas whose the life stage most likely to benefit from camouflage, show similar effects under ecologically realistic viewing contexts.

A further important consideration is that juvenile pandas may be viewed against different background types depending on their behaviour and position in the habitat. When on the ground, they are likely to be seen against soil, rocks, leaf litter and low vegetation. When climbing or resting in trees, they may be viewed against canopy backgrounds composed of branches, foliage, shaded regions and bright sky gaps. These canopy scenes can contain strong luminance contrasts and complex edges (Frazor and Geisler, 2006), which may better correspond to the black-and-white fur. Additionally, vegetative cover can partly obscure the animal’s outline, making it harder for predators to detect or recognize targets (DiPietro et al., 2002, Sovrano and Bisazza, 2008). Experimental studies using predator models such as sparrowhawks (Tvardíková and Fuchs, 2010a) and leopards (Coss et al., 2005) show that prey animals respond more slowly to threats partially hidden by vegetation, indicating that such occlusion impedes target detection. For juvenile pandas, partial cover from branches and leaves may therefore interact with pelage patterning to influence camouflage-relevant visual metrics.

We used artificial panda models with natural and experimentally modified colour patterns, to test whether juvenile panda colouration produces visual effects consistent with background matching and/or disruptive coloration, and whether these effects depend on environmental context. Specifically, we asked whether natural juvenile panda colouration reduces target-background contrast relative to altered colour or brightness treatments, whether the natural arrangement of black, white and grey regions produces stronger edge-disruption metrics than modified pattern arrangements, and whether these camouflage-relevant effects are influenced by background position, vegetation occlusion and illumination. We predicted that naturally patterned models would show lower colour and/or luminance contrast against woodland backgrounds than models with altered brightness or colour composition, and higher edge-disruption metrics than models with modified pelage arrangements. We further predicted that partial vegetation occlusion would reduce contrast and increase edge disruption, and that canopy backgrounds would provide a more favourable visual context for camouflage than ground backgrounds.

## Methods

### Model design and experimental treatments

The juvenile panda models were designed to be the lateral view approximate dimensions of a six-month-old juvenile panda (70 cm wide × 43 cm tall, Schaller, 1985). Models were cut from 6 mm thick birch plywood (The Grain Ltd. Liskeard, UK), leaving a sharp stake about 5 cm long at the hinder limb for insertion into the ground. The size of the models varied slightly among replicas. Lateral images containing the different experimental patterns were printed on matte paper and stuck to the plywood boards. The pattern manipulation of the brightness, pattern patch and margin interaction, and colour reverse were inspired by Cuthill et al. (2005), Stevens et al. (2006) and Honma et al. (2015).

The natural coloration (NM) was used as the reference treatment. Modified treatments altered patch extent, overall luminance, internal contrast, the positions at which pattern boundaries intersected the body outline, or the presence of simulated vegetation shadows; an additional ground-position treatment (GD) used the naturally patterned model to compare canopy and ground-level backgrounds; its placement protocol is described below (Figure 2, Table 1).

**Table 1.**
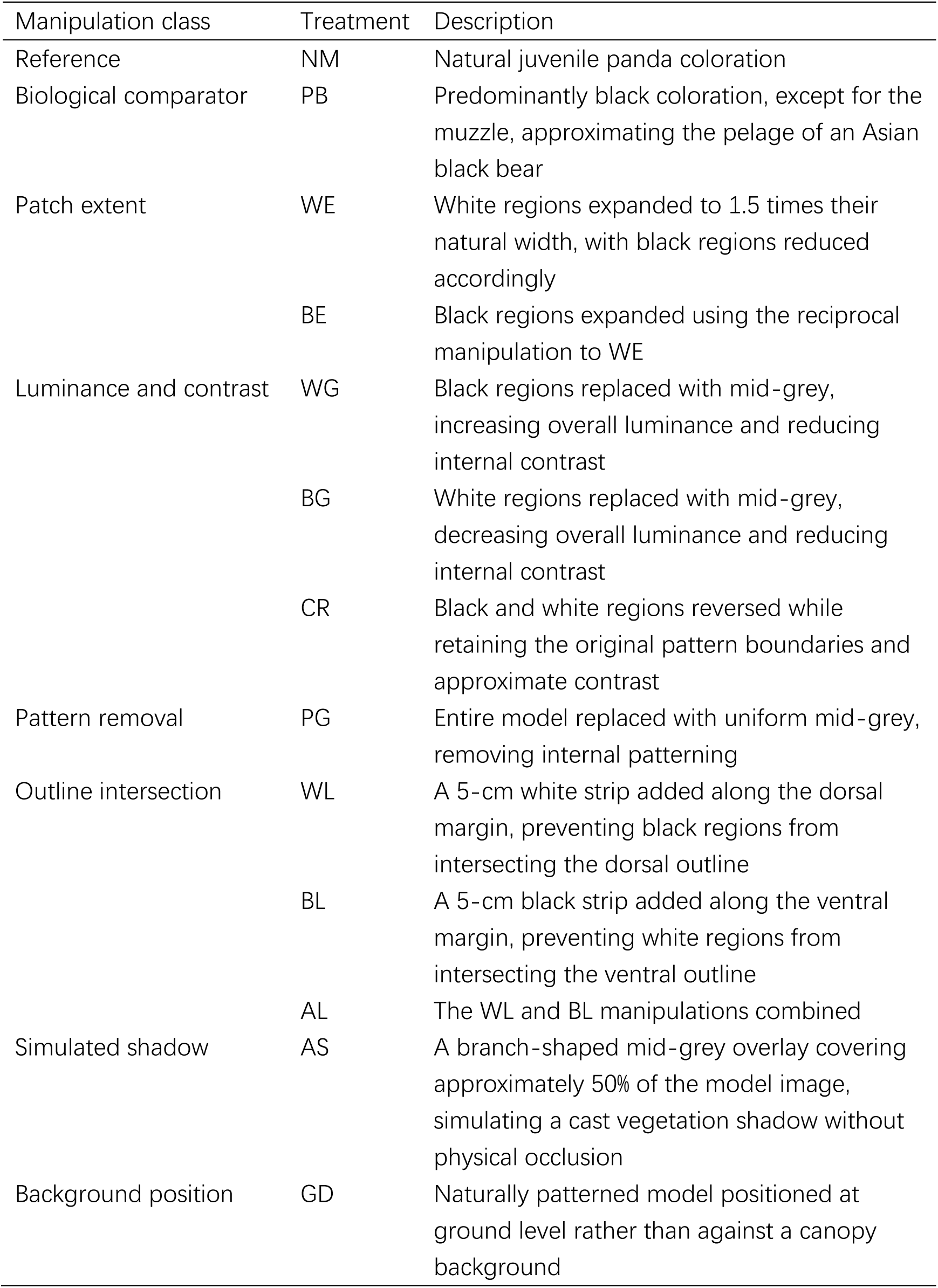
Experimental treatments applied to juvenile giant panda models.

Background-matching analyses compared NM with PB, WE, BE, WG, BG and PG. Edge-disruption analyses compared NM with CR, WG, BG, WL, BL, AL and PG. AS was used to distinguish the visual effects of simulated cast shadows from those of physical vegetation occlusion, whereas GD was used to compare camouflage-relevant metrics between canopy and ground-level backgrounds. Further details of treatment construction and placement are provided in the Supplementary Methods.

### Model placement and photography

The study was conducted between February and July 2022 at College Reservoir (50°9’37 N, 5°7’22 W) of Penryn in Cornwall, UK. Twenty trees meeting predefined suitability criteria were selected for photography: taller than 3 m, had accessible branches below 3 m, and had shrubs growing beneath. The maximum placement height was limited by the length of telescopic rod. To simulate juvenile pandas hiding in trees, models were placed on a branch near the trunk, raised by a 3-meter-long aluminium alloy telescopic rod. From the camera position, the background consisted of branches, foliage and gaps of visible sky (Figure 1b).

**Figure 1.**
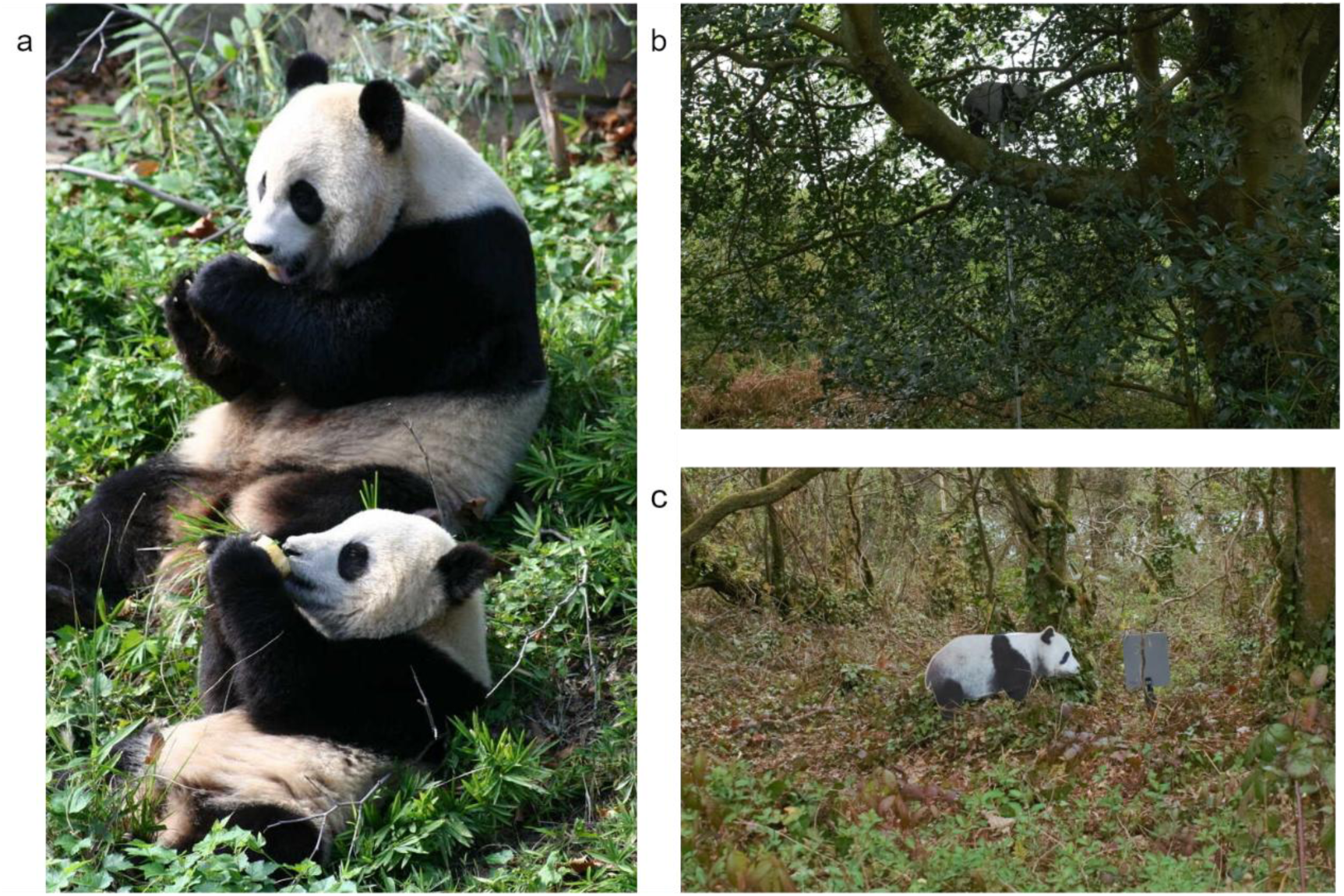
a. Adult and juvenile giant panda; b. Panda model positioned against a canopy background and partially obscured by branches; c. Panda model positioned on a ground-level background without vegetation obstruction.

**Figure 2.**
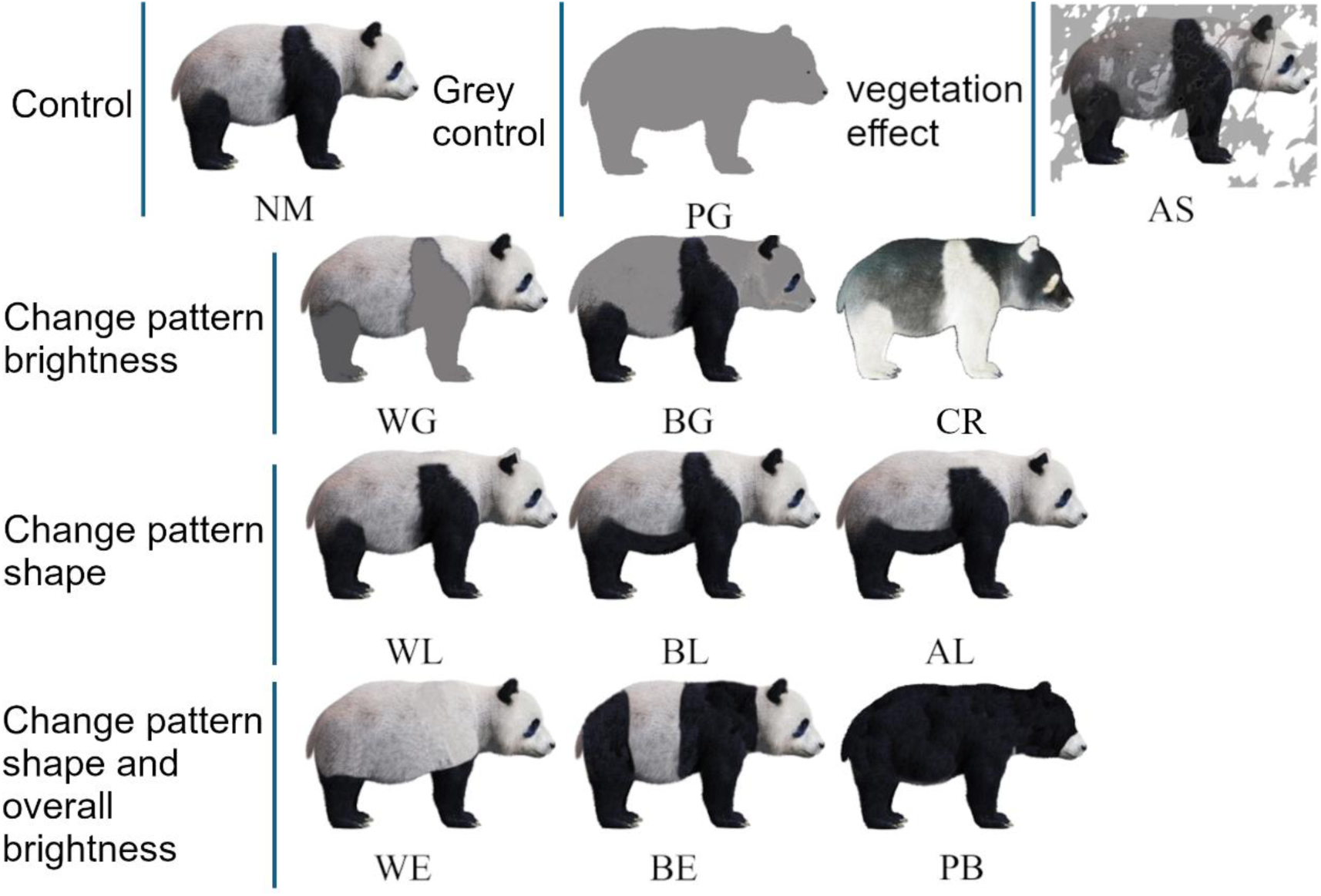
Colour and pattern treatments applied to juvenile giant panda models. NM retained the natural juvenile panda coloration and served as the reference treatment, whereas PG was uniformly mid-grey and matched the approximate mean luminance of NM. Experimental treatments altered: (1) patch luminance or luminance polarity while retaining the principal pattern boundaries: WG, black regions replaced with grey; BG, white regions replaced with grey; and CR, black and white regions reversed; (2) the positions at which pattern boundaries intersected the body outline: WL, white regions linked along the dorsal margin; BL, black regions linked along the ventral margin; and AL, the WL and BL manipulations combined; (3) patch extent and overall luminance: WE, white regions expanded; BE, black regions expanded; and PB, predominantly black coloration; and (4) simulated vegetation shadowing: AS, a branch-shaped grey overlay covering approximately 50% of the model image.

To compare canopy and ground-level background, the panda models with natural coloration were photographed on the ground level and positioned in the background of shrubs and grass(Figure 1c), coded as the ground treatment (GD).

Vegetation occlusion was manipulated at both canopy and ground-level positions. In the behind vegetation condition (B), panda models were put behind the branches, leaves, or shrub. In the front-of-vegetation condition (F), the model was positioned between the camera and the vegetation and was therefore not physically occluded.

Because the shadows cast by the branches on the model under direct sunlight may have created additional ‘patterns’ which may affect camouflage, the experiment was carried out on sunny days when strong shadows were cast by the trees, and cloudy days when no strong shadows were cast on the panda model. When photographed under direct sunlight, the camera was avoided directly facing into or from behind the sun, to reduce the risk of overexposure or excessive reflection from the model.

For photographing the panda models, a Sony A7 digital camera with a 28-70mm lens was used, positioned approximately 3-5 meters away from the model under natural light. Manual adjustments were made to the camera settings for white balance, setting the aperture at f/6.3 and ISO at 160. A tripod was used to stabilise the camera and to ensure that every photo was taken at the same height. A photography umbrella was used to prevent glare, and a DIGITAL HD ® MENNON 18% grey standard reflectance scale bar was included in each image for calibration (Stevens et al., 2007, Troscianko and Stevens, 2015). All photos were taken in RAW format. In total, 1120 images were captured during the experiment.

### Image analysis and visual modelling

All image processing and analyses were performed using the Multispectral Image Calibration and Analysis Toolbox v2.2.2 (Troscianko and Stevens, 2015) in ImageJ 1.53v (Schneider et al., 2012). The panda model outline in each image was manually selected as the region of interest (ROI). The background was defined as the circular area with a radius twice that of the boundary circle, centred on the panda image. This larger background area was necessary because the panda model occupied most of the space within the original boundary circle, leaving insufficient area to accurately represent the surrounding environment. The images underwent calibration using the grey standard to normalize the light conditions and were saved as 32-bit multispectral images in.tiff format (Troscianko and Stevens, 2015). Subsequently, a scale bar was added in accordance with the length of the panda model in each photo. For each photo, we converted the data into the cone catch images of the domestic dog (Neitz et al., 1989) and domestic cat (Loop et al., 1987) which represent the potential feline and canine predators’ vision.

Acuity-corrected (Caves and Johnsen, 2018) views with distance scaling were applied to the image analysis to account for the distance-dependent blur, using domestic dog to represent the canine predator vision and visual acuity as 12 cycles per degree (cpd) (Odom et al., 1983), and cat as feline predators and visual acuity as 9 cpd (Hall and Mitchell, 1991). Images were modelled at viewing distances of 5, 10, 20, 30 and 40 m.

### Target–background contrast

We assessed the colour congruence level between the panda model and their backgrounds with the logarithmic form of the receptor noise model (RNL) (Vorobyev and Osorio, 1998). Chromatic contrast was expressed as ΔS and luminance contrast as ΔS_lum_, both in units of just-noticeable difference (JND). Values below 1 JND is considered indistinguishable under ideal viewing conditions, while increasing values above this suggest increasingly discernible differences (Siddiqi et al., 2004, Stevens et al., 2017, Perez-Martinez et al., 2020). Receptor noise parameterised by the Weber fractions (ω) for the domestic dog chromatic vision was 0.05 with cone ratio L/M:S=18820:1842 (Mowat et al., 2008); and for cat chromatic vision was Weber fraction of 0.05 and cone ratio L/M:S=6:1 (Clark and Clark, 2016). The Weber fraction of luminance channel was set as 0.22 for dog (Pretterer et al., 2004) and 0.20 for cat (Clark and Clark, 2016). It should be noted that pandas are essentially achromatic, so colour analysis should be interpreted with caution since small differences in colour in otherwise white, grey, or black markings may produce exaggerated discrimination values.

### Edge-disruption

We used the Gabor filtering to quantify achromatic edge disruption, which measures the proportion of false edges orthogonal to the animal’s true outline (Troscianko et al., 2017). Since luminance contrast and edge orientation changes are often the main factors affecting disruptive coloration, the analysis focused only on the luminance channel. The GabRat sigma value was set at 3. The animal vision and distance scaling was applied when performing the GabRat analysis to investigate if the effect varied with viewing distance. Following Troscianko et al. (2017), higher GabRat values suggest greater pattern-outline interference, values below 0.2 indicate potentially low-level edge disruption and above 0.4 indicates potentially high-level disruption. It should be noted, however, larger GabRat measures arise from high contrast, which is also common with conspicuous markings. Therefore, its interpretation as evidence for camouflage in this study relies on the experimental manipulations that follow.

### Statistical Analysis

Initial examination of the ΔS of colour and ΔS_lum_ values revealed a positive skew, prompting the application cube root transformation on ΔS values for normalization. A linear mixed-effect model (LMM) fitted with restricted maximum likelihood was fitted for each response variable. Fixed effects in the models included pattern (i.e. treatment), weather condition (sunny / cloudy) and vegetation occlusion (behind or in front of vegetation), and the interactions between them. Site ID was included as random effects, mitigating the influence of site selection and canopy density conditions.

response ∼ Pattern + Weather + Vegetation + Pattern:Weather + Pattern:Vegetation + Weather:Vegetation + (1 | Site)

An ANOVA was employed to determine the significance of effects or interactions, followed by Tukey post hoc tests for within-effect level comparisons. If the interaction was significant, the treatments would be compared separately on each moderator variable level. All statistical tests were conducted in R v4.2.3 (R Core Team, 2021) using the lme4 package v1.1-35.1 (Bates et al., 2015) and emmeans package (Lenth, 2023).

## Results

### Edge disruption

Based on the image analysis, we found that the panda models exhibit a relatively high edge disruption level against the natural background, and that most of the treatments had a GabRat value over 0.3, suggests the high pattern–outline interference in panda coloration (Figure 3). The LMM model showed that the interaction between pattern and weather (dog: F = 3.258, p<0.001; cat: F = 1.703, p=0.043), as well as the interaction between pattern and vegetation (dog: F = 9.086, p<0.001; cat: F = 5.256, p<0.001), had a significant effect on the GabRat value, whereas the interaction between weather and vegetation had no significant effect (Table 2), indicating that the effect of pattern on potential edge disruption may differ across light conditions (cloudy or sunny) and vegetation occlusion (behind or in front of vegetation).

**Figure 3.**
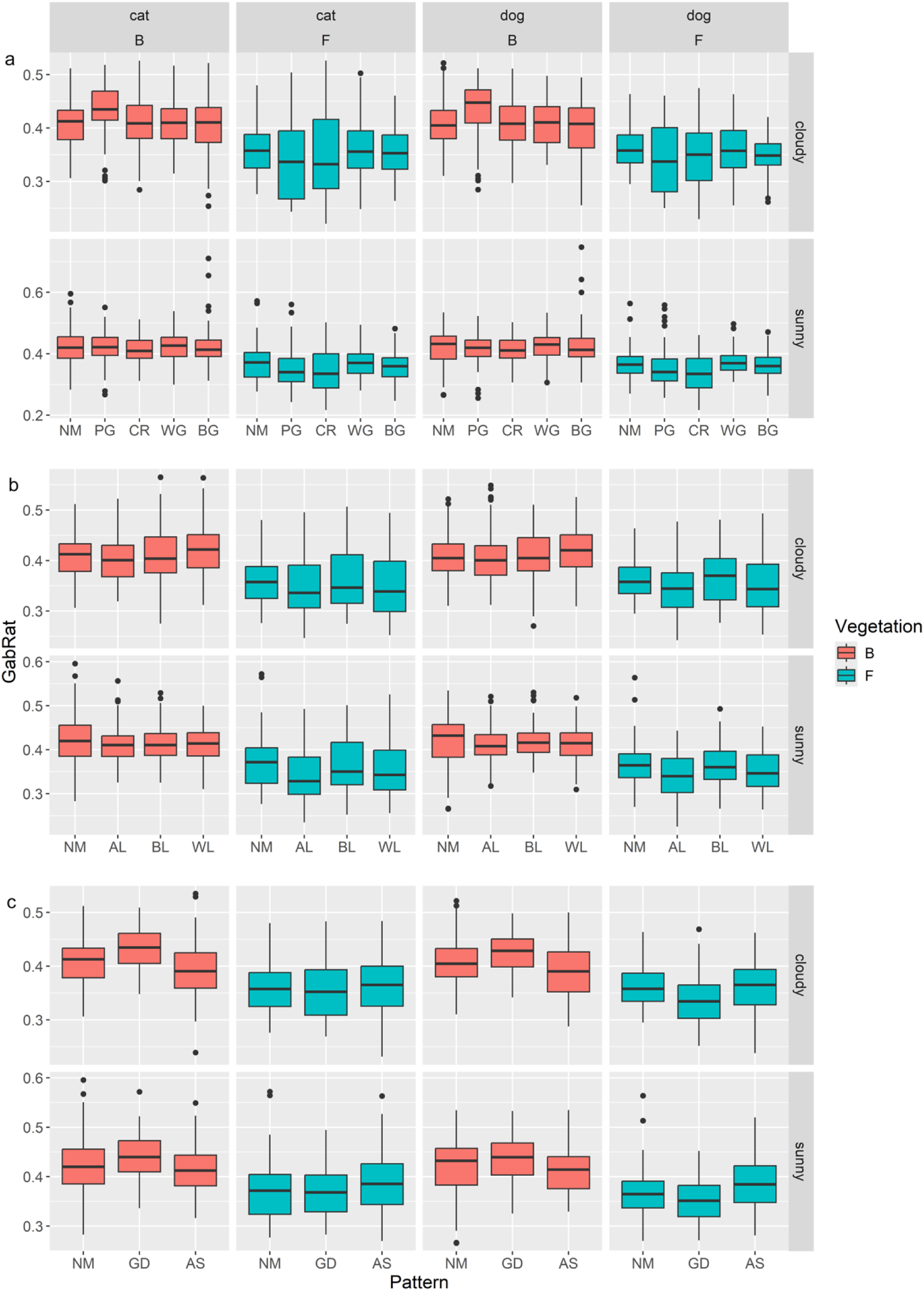
GabRat values for panda pattern treatments viewed under domestic cat and dog vision. Results are shown for cloudy and sunny conditions, with vegetation occlusion (B) and no occlusion (F). Panels compare natural coloration with treatments that: (a) alter colour, (b) alter pattern shape, and (c) manipulate cast shadow or model height. Boxplots show the median and 25th and 75th percentiles; the whiskers indicate the values within 1.5 times the interquartile range.

**Figure 4.**
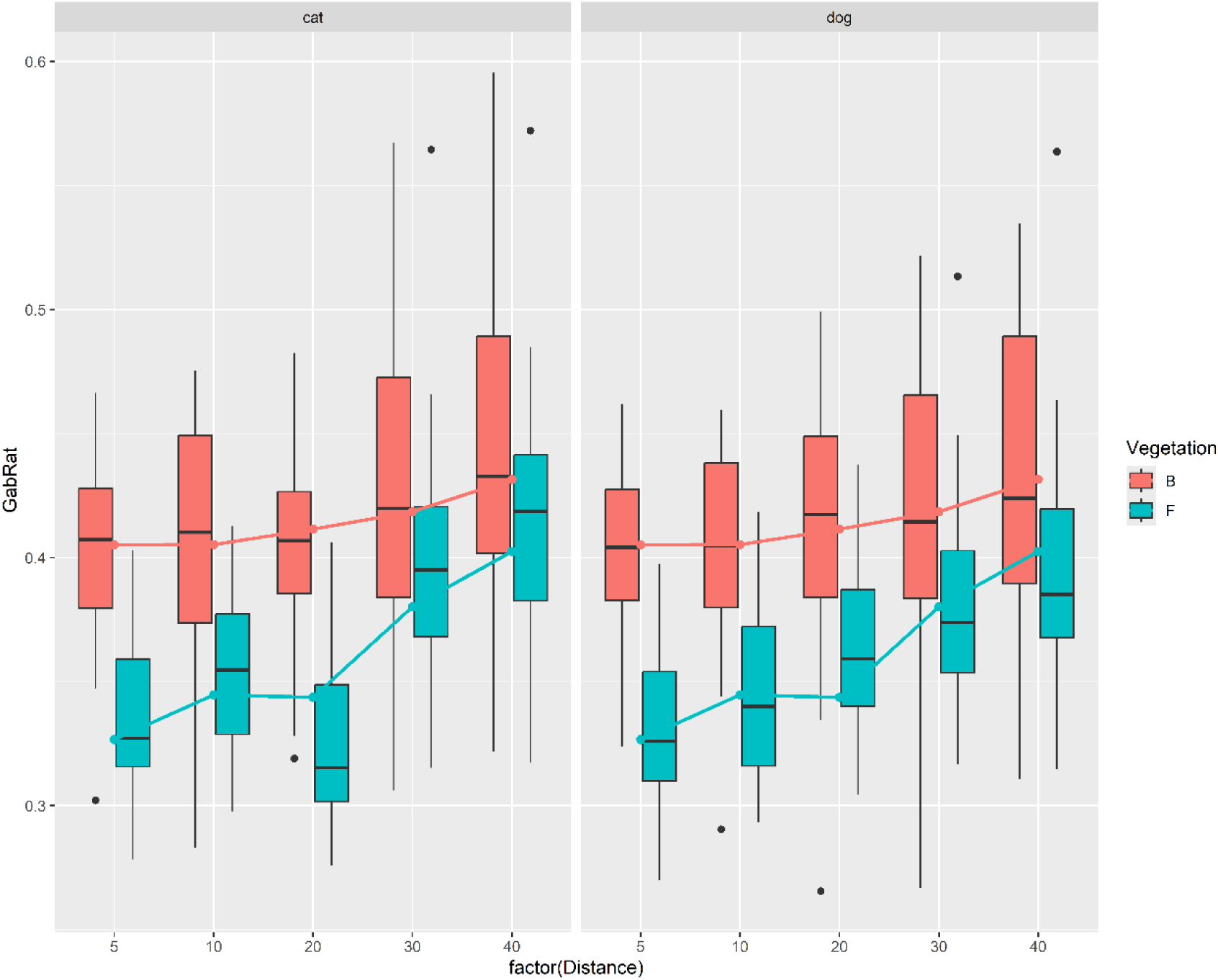
The edge disruption of panda models under domestic dog and cat vision at 5-40m distance under two vegetation conditions (B: covered by vegetation/F: without vegetation occlusion). The box plots show the median and 25th and 75th percentiles; the whiskers indicate the values within 1.5 times the interquartile range.

**Table 2.** Summary of results comparing of influence factors for all camouflage metrics.

| Species | Metrix | Fixed effect | Mean Sq | F value | Pr(>F) |
| --- | --- | --- | --- | --- | --- |
| Dog | GabRat | Pattern | 0.011 | 4.666 (9, 3329.4) | <0.001 |
|  |  | Weather | 0.059 | 25.826 (1, 3331.5) | <0.001 |
|  |  | Vegetation | 2.726 | 1197.286 (1, 3327.1) | <0.001 |
|  |  | Pattern:Weather | 0.007 | 3.258 (9, 3328.8) | 0.001 |
|  |  | Pattern:Vegetation | 0.021 | 9.086 (9, 3327.1) | <0.001 |
|  |  | Weather:Vegetation | 0.004 | 1.805 (1, 3327.1) | 0.179 |
|  | dS | Pattern | 0.729 | 2.77 (8, 552.8) | 0.005 |
|  |  | Weather | 0.234 | 0.888 (1, 553.8) | 0.346 |
|  |  | Vegetation | 184.571 | 701.414 (1, 552) | <0.001 |
|  |  | Pattern:Weather | 0.087 | 0.332 (8, 552.6) | 0.954 |
|  |  | Pattern:Vegetation | 0.824 | 3.131 (8, 552) | 0.002 |
|  |  | Weather:Vegetation | 0.061 | 0.233 (1, 552) | 0.629 |
|  | dS (Luminance) | Pattern | 1.002 | 14.796 (8, 552.7) | <0.001 |
|  |  | Weather | 0.051 | 0.753 (1, 553.5) | 0.386 |
|  |  | Vegetation | 0.516 | 7.622 (1, 552) | 0.006 |
|  |  | Pattern:Weather | 0.032 | 0.472 (8, 552.5) | 0.876 |
|  |  | Pattern:Vegetation | 0.101 | 1.496 (8, 552) | 0.156 |
|  |  | Weather:Vegetation | 0.001 | 0.018 (1, 552) | 0.892 |
| Cat | GabRat | Pattern | 0.015 | 5.352 (9, 3330.3) | <0.001 |
|  |  | Weather | 0.038 | 13.717 (1, 3333.4) | <0.001 |
|  |  | Vegetation | 2.663 | 957.09 (1, 3327.1) | <0.001 |
|  |  | Pattern:Weather | 0.005 | 1.703 (9, 3329.3) | 0.043 |
|  |  | Pattern:Vegetation | 0.015 | 5.256 (9, 3327.1) | <0.001 |
|  |  | Weather:Vegetation | 0.001 | 0.512 (1, 3327.1) | 0.474 |
|  | dS | Pattern | 0.253 | 2.724 (8, 552.8) | 0.006 |
|  |  | Weather | 0.024 | 0.255 (1, 553.7) | 0.614 |
|  |  | Vegetation | 95.827 | 1030.036 (1, 552) | <0.001 |
|  |  | Pattern:Weather | 0.020 | 0.214 (8, 552.5) | 0.988 |
|  |  | Pattern:Vegetation | 0.349 | 3.748 (8, 552) | <0.001 |
|  |  | Weather:Vegetation | 0.033 | 0.352 (1, 552) | 0.553 |
|  | dS (Luminance) | Pattern | 1.081 | 14.979 (8, 552.7) | <0.001 |
|  |  | Weather | 0.048 | 0.664 (1, 553.5) | 0.416 |
|  |  | Vegetation | 0.545 | 7.557 (1, 552) | 0.006 |
|  |  | Pattern:Weather | 0.034 | 0.473 (8, 552.5) | 0.875 |
|  |  | Pattern:Vegetation | 0.109 | 1.516 (8, 552) | 0.149 |
|  |  | Weather:Vegetation | 0.001 | 0.011 (1, 552) | 0.917 |

Pairwise comparison results showed that the reduction of the pattern-outline interaction (AL - NM) had a negative effect on GabRat, when the models in the absence of vegetation occlusion and under sunny weather in both dog and cat vision (Table S1, S2). However, no significant difference was found when the intersection between the body outline and either the single black or white patches was removed. By contrast, the colour reverse (CR) treatment decreased GabRat values compared with the control when the models were under the same conditions (Table S1, S2), suggesting that the natural arrangement of both black and white patches is important for generating this high GabRat effect. Treatments designed to reduce the contrast between black and white patches (WG and BG) did not significantly differ from the control treatment in GabRat. The only significant difference occurred in the absence of vegetation under cat vision, where the uniformly grey treatment had a lower GabRat than the natural pattern (PG - NM (F): cat estimate = - 0.019, p=0.043).

In addition, the panda models under sunny weather showed slightly but significantly higher GabRat than under cloudy weather (cloudy - sunny: dog estimate =-0.009, p<0.001, cat estimate =-0.007, p<0.001). The placement of the model behind vegetation also significantly increased the GabRat of targets (B - F: dog estimate = 0.056, p<0.001, cat estimate = 0.055, p<0.001) (Figure 3).

The GabRat changes with distance showed that the potential edge disruption of naturally patterned juvenile pandas increases with distance. Panda models hidden in the branches (B) showed a high edge disruption level (>0.4) even at the minimum observation distance, higher than those exposed in front of branches (F). The difference between vegetation conditions narrowed with increasing viewing distance, with GabRat values in the unobstructed condition approaching those under vegetation occlusion by 40 m.

### Target–background contrast

Pattern treatment and vegetation significantly affected the chromatic contrast and luminance contrast, but the lighting environment (i.e. sunny/cloudy) had no effect (Table 2). Only the interaction between pattern and the vegetation had a significant effect on colour contrast (dog: F = 3.131, p=0.002; cat: F = 3.748, p<0.001), the interactions of weather with patterns or vegetation did not significantly affect the background matching.

Specifically, expanding the black pattern or white pattern increased the colour contrast between the panda model and the canopy background in both the dog and cat visual models when the model is under the cover of vegetation occlusion (BE-NM: dog estimate = 0.456, p=0.004, cat estimate = 0.308, p=0.001; WE-NM: dog estimate = 0.454, p=0.004, cat estimate = 0.347, p<0.001). In contrast, replacing either the black or white regions with grey, as in the WG and BG treatments, had no significant effect. Moreover, the natural panda coloration (NM) did not show any advantage in colour matching compared to either the black (PB) or grey (PG) treatments (Table S3, Figure 5).

**Figure 5.**
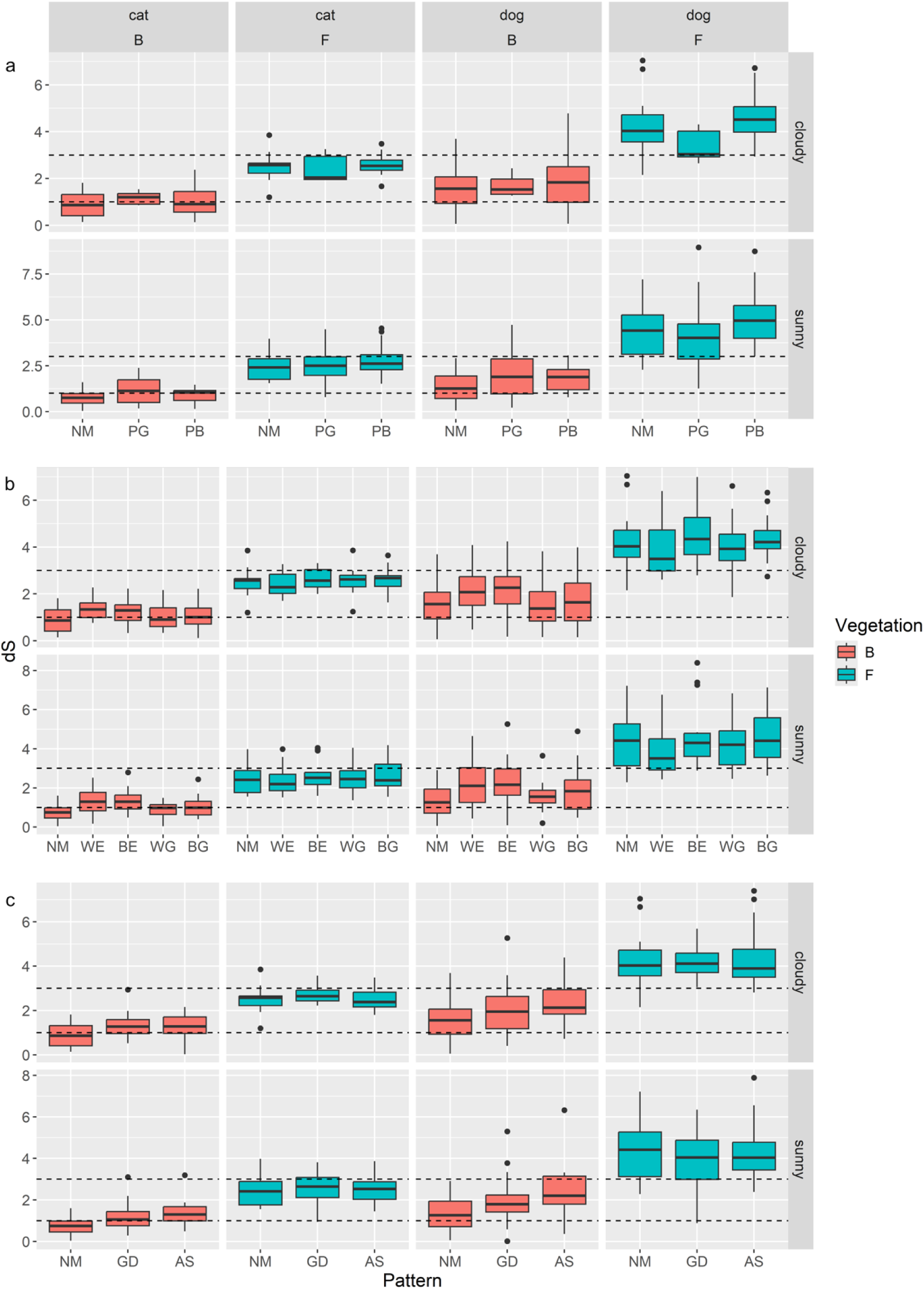
The colour contrast for panda pattern treatments viewed under domestic cat and dog vision. Results are shown for cloudy and sunny conditions, with vegetation occlusion (B) and no occlusion (F). Panels compare natural coloration with treatments that: (a) alter colour, (b) alter pattern brightness, and (c) manipulate cast shadow or model height. Boxplots show the median and 25th and 75th percentiles; the whiskers indicate the values within 1.5 times the interquartile range. The dotted line represents 1 and 3 JND.

The luminance contrast was not affected by the pattern expansion and brightness change of the model, whereas the black bear treatment showed a significantly higher contrast than the control group (PB-NM: dog estimate = 0.276, p<0.001, cat estimate = 0.287, p<0.001). The vegetation obstacle also significantly decreased the luminance contrast (B-F: dog estimate =-0.059, p=0.006, cat estimate =-0.061, p=0.006) (Table S4, Figure 6).

**Figure 6.**
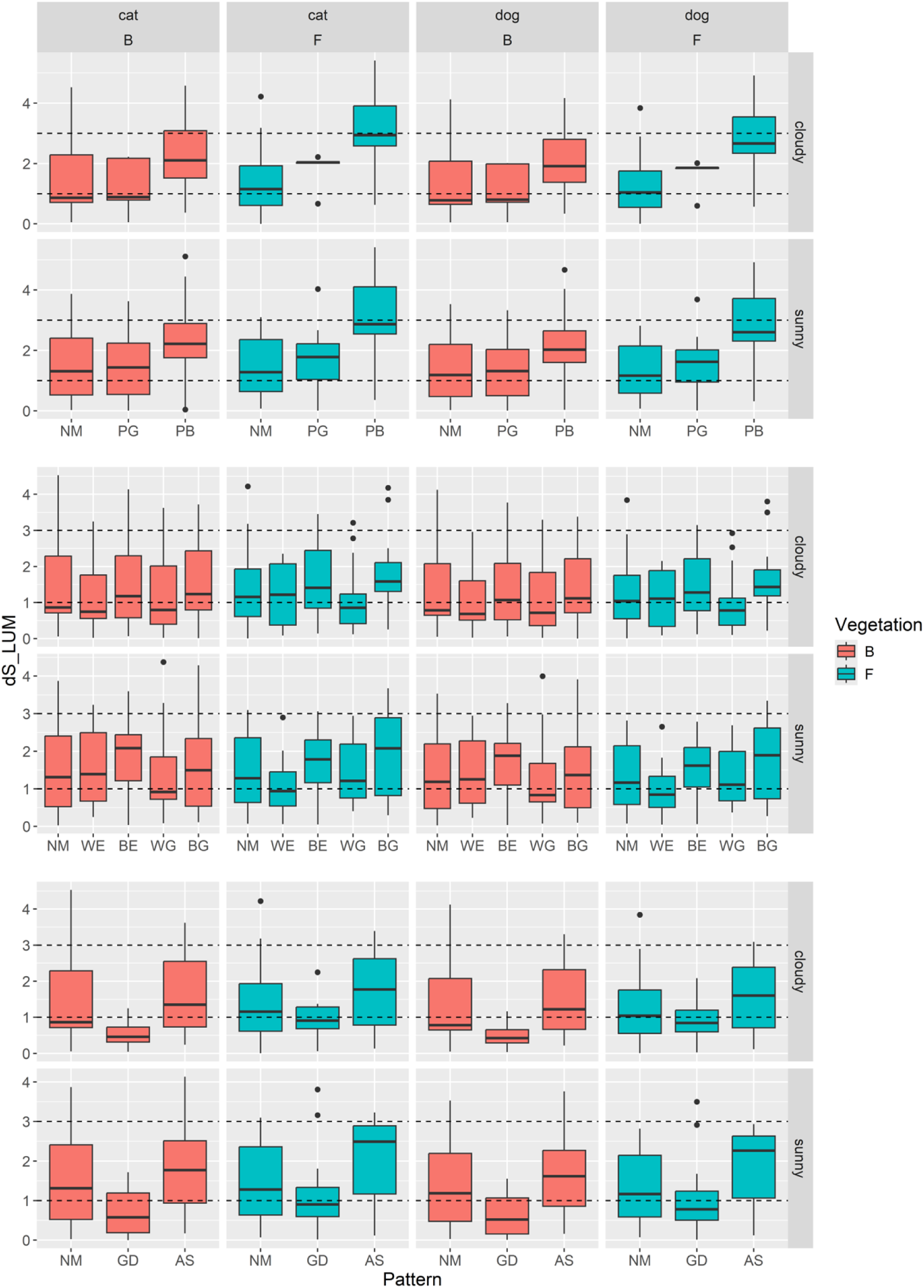
The luminance contrast for panda pattern treatments viewed under domestic cat and dog vision. Results are shown for cloudy and sunny conditions, with vegetation occlusion (B) and no occlusion (F). Panels compare natural coloration with treatments that: (a) alter colour, (b) alter pattern brightness, and (c) manipulate cast shadow or model height. Boxplots show the median and 25th and 75th percentiles; the whiskers indicate the values within 1.5 times the interquartile range. The dotted line represents 1 and 3 JND.

### The effect of model height and shadow

The comparison between the canopy background and the ground showed that when the panda model was photographed on the ground, it had lower luminance contrast in both kinds of visual system (GD-NM: dog estimate = - 0.175, p=0.005, cat estimate =-0.181, p = 0.005) (Table S4, Figure 6). However, under vegetation occlusion, models positioned at ground level had higher chromatic contrast than those viewed against canopy backgrounds under the cat visual model (GD-NM estimate = 0.318, p = 0.001). The corresponding estimate under the dog visual model was no significant difference (estimate = 0.348, p = 0.101), or in both visual systems when vegetation was absent (Table S3, Figure 5). Moreover, against a ground background GabRat values were higher than against a canopy background under vegetation (GD-NM B: dog estimate = 0.016, p = 0.030; cat estimate = 0.018, p = 0.017). In the absence of vegetation occlusion, however, the ground-level treatment had lower GabRat than the canopy treatment under the dog visual model (estimate =-0.017, p = 0.009) whereas no significant difference was detected under the cat vision (estimate =-0.003, p = 1.000; Table S1, Figure 3).

These results suggest that the effect of model position on the measurements of GabRat is mediated by both background type and viewer visual system. Under vegetation occlusion, canopy-positioned models had lower chromatic contrast than ground-level models under the cat visual model, with a similar but non-significant direction under the dog visual model. Ground-level positioning reduced luminance contrast under both visual systems, whereas the effect of model position on edge disruption depended on both vegetation occlusion and the observer visual model.

The artificial-shadow treatment had no significant difference from the control on any camouflage metric to either animal’s visual model (Table S1-S3). Artificially adding a branch-shaped grey shadow therefore produced no detectable change in the camouflage-relevant visual metrics measured here.

## Discussion

We employed quantitative methods to test whether the giant panda’s distinctive black-and-white fur potentially contributes to camouflage through background matching and disruptive coloration. The results suggest that juvenile panda coloration could produce camouflage-related visual effects, influencing both target-background contrast and edge-disruption metrics, but the strength and direction of these effects depended on the specific treatment comparison, background, vegetation condition and visual model. The spatial extent and arrangement of the black and white patches were more influential than moderate changes in the contrast between them. By incorporating vegetation and the shadows this casts into the analysis, we found that pandas in dense vegetation exhibited lower contrast and greater predicted edge disruption, whereas artificial cast shadows had no detectable effect. Furthermore, lighting conditions influenced only the effectiveness of putative disruptive coloration, with direct light having an interaction with panda coloration and the vegetation in the background, likely making detection more difficult. Comparisons between models placed on the ground or in 3m high in tree canopy revealed that pandas have lower colour contrast in the tree canopy compared to the ground to felid vision. These findings suggest that the juvenile panda coloration can interact with the structural and illumination properties of woodland habitats to generate several complementary camouflage-relevant visual effects, which may relevant to tree-climbing behaviour and felid-like visual systems, and operate in concert with other anti-predator strategies.

Previous research has suggested that adult panda coloration may facilitate both background matching and edge disruption, particularly when viewed at relatively long distances (Caro et al., 2017, Nokelainen et al., 2021). Our study extends this work by focusing on body size in juveniles, which represent the life stage most vulnerable to predation. The panda’s GabRat values were already relatively high at distances as close as 5 meters when models partially obscured by vegetation. In combination with a reduction in contrast through background matching, this could potentially further reduce detection risk at close range. As body size increases, the mismatched visual surface area expands, reducing camouflage efficiency (Remmel and Tammaru, 2009, Skelhorn et al., 2010b, Pembury Smith and Ruxton, 2021). Thus, although adult panda coloration may still serve a camouflaging role, its effectiveness diminishes, and adults may rely less on camouflage due to the strengthening of secondary defences (Stevens and Ruxton, 2012). Accordingly, studies on anti-predator strategies in juvenile pandas may more accurately reflect the species’ ecological context.

Altering the contrast of the panda’s pelage did not significantly affect measurements of predicted edge disruption, suggesting that changes in coloration, for example due to dirt from the environment, have minimal effects. Alternatively, the introduction of intermediate colours may compensate for contrast loss by enabling effective matching with background elements (Caro et al., 2017, Nokelainen et al., 2021). Some supporting evidence comes from studies on owls, where grey morphs are less detectable against snowy and bark backgrounds compared to brown morphs (Koskenpato et al., 2020), and from research indicating that dirt-covered avian feathers enhance camouflage (Delhey et al., 2007, Ruxton and Stevens, 2015).

Natural panda models had better potential camouflage in the canopy compared with on the ground under felid vision when compared to canid vision. These findings suggest that panda coloration may have evolved as an anti-predator strategy specifically adapted to felid visual systems by reducing the colour contrast in canopy background. In evolutionary arms races, animal coloration may develop camouflage targeted toward specific visual systems of observers, making it more effective against those observers even if it remains conspicuous to others. For example, the tiger’s coloration blends effectively with green vegetation under dichromatic vision (Fennell et al., 2019). Given the predation pressure juvenile pandas face from leopards (Schaller, 1985), such an evolutionary direction likely benefits individual survival. The panda’s white fur may be an adaptation for camouflage against the bright sky. Research on the white colour morph of the black bear (*Ursus americanus kermodei*) in British Columbia has shown that the white morph is associated with reduced detection against the light background of the sky when viewed from a lower angle by fish (Klinka and Reimchen, 2009, Reimchen et al., 2021), which to some extent supports the present experimental results.

Canopy-level backgrounds may facilitate juvenile pandas’ camouflage, aligning with their tree-climbing behaviour used to evade threats, which suggests that juveniles may behaviourally select suitable backgrounds to enhance camouflage. By occupying visually complex or partially obscured locations, juveniles may reduce the visibility of body contours and other diagnostic features, thereby delaying detection or recognition (DiPietro et al., 2002, Coss et al., 2005, Tvardíková and Fuchs, 2010b, Dimitrova and Merilaita, 2009). Moreover, animals may enhance their camouflage by adjusting their posture (Barbosa et al., 2012, Suzuki and Sakurai, 2015, McEwen et al., 2024). In this study, all panda models were posed standing upright on the ground; however, juvenile pandas in the wild may adopt seated or climbing postures, which could alter the appearance and any disruptive effect of their body patterns due to changes in limb placement and body orientation to improve concealment. However, which plant species provide more effective concealment and how concealment selection can optimise camouflage remain areas for further investigation.

This research was conducted in a non-native environment in the UK, rather than in natural habitats of pandas, where vegetation type, background structure, and illumination are different. the results provide quantitative evidence that juvenile panda coloration can produce visual effects consistent with background matching and disruptive coloration, and highlight how coat pattern, habitat structure, predator vision and behaviour may interact to influence camouflage. Future studies should aim to test these findings in situ within the panda’s natural habitat to better understand how these camouflage mechanisms function under ecologically realistic conditions.

## Conflict of Interest

The authors declare no conflicts of interest.

## Author contributions

**Zixiang Li:** Conceptualization; Methodology; Investigation; Data curation; Formal analysis; Software; Visualization; Funding acquisition; Writing – original draft; Writing – review & editing.

**Laura A. Kelley:** Methodology; Supervision; Writing – review & editing.

**Martin Stevens:** Conceptualization; Methodology; Supervision; Project administration; Writing – review & editing.

## Notes

### Competing Interest Statement

The authors have declared no competing interest.

### Funding

- China Scholarship Council: 202006040047

