## Supplementary material for "Camouflage functions of juvenile giant panda colouration in forest environments": Table S1

| **Edge disruption effect pairwise comparison between the pattern treatments on different vegetation situation level.** | | | | | | | |
| --- | --- | --- | --- | --- | --- | --- | --- |
| contrast | Vegetation | dog | | | cat | | |
|  |  | estimate | SE | p.value | estimate | SE | p.value |
| PG-NM | B | 0.002 | 0.006 | 1.000 | 0.001 | 0.007 | 1.000 |
| AL-NM |  | -0.007 | 0.005 | 0.829 | -0.009 | 0.006 | 0.697 |
| BL-NM |  | -0.002 | 0.005 | 1.000 | -0.003 | 0.006 | 0.999 |
| WL-NM |  | 0.000 | 0.005 | 1.000 | -0.002 | 0.006 | 1.000 |
| BG-NM |  | -0.005 | 0.005 | 0.986 | -0.004 | 0.006 | 0.999 |
| WG-NM |  | 0.003 | 0.005 | 0.999 | 0.000 | 0.006 | 1.000 |
| CR-NM |  | -0.004 | 0.005 | 0.990 | -0.006 | 0.006 | 0.944 |
| GD-NM |  | 0.016 | 0.005 | **0.030** | 0.018 | 0.006 | **0.017** |
| AS-NM |  | -0.014 | 0.005 | 0.051 | -0.015 | 0.006 | 0.063 |
| PG-NM | F | -0.013 | 0.006 | 0.259 | -0.019 | 0.007 | **0.043** |
| AL-NM |  | -0.020 | 0.005 | **0.001** | -0.021 | 0.006 | **0.002** |
| BL-NM |  | 0.003 | 0.005 | 1.000 | -0.001 | 0.006 | 1.000 |
| WL-NM |  | -0.013 | 0.005 | 0.112 | -0.015 | 0.006 | 0.072 |
| BG-NM |  | -0.009 | 0.005 | 0.587 | -0.008 | 0.006 | 0.761 |
| WG-NM |  | 0.003 | 0.005 | 1.000 | -0.001 | 0.006 | 1.000 |
| CR-NM |  | -0.021 | 0.005 | **0.001** | -0.020 | 0.006 | **0.005** |
| GD-NM |  | -0.017 | 0.005 | **0.009** | -0.003 | 0.006 | **1.000** |
| AS-NM |  | 0.011 | 0.005 | 0.229 | 0.008 | 0.006 | 0.750 |

Table S2

| **Edge disruption effect pairwise comparison between the pattern treatments on different weather situation level.** | | | | | | | |
| --- | --- | --- | --- | --- | --- | --- | --- |
| contrast | Weather | dog | | | cat | | |
|  |  | estimate | SE | p.value | estimate | SE | p.value |
| PG-NM | cloudy | -0.002 | 0.008 | 1.000 | -0.005 | 0.009 | 0.999 |
| AL-NM |  | -0.008 | 0.005 | 0.706 | -0.008 | 0.006 | 0.770 |
| BL-NM |  | 0.003 | 0.005 | 0.998 | 0.002 | 0.006 | 1.000 |
| WL-NM |  | 0.000 | 0.005 | 1.000 | -0.001 | 0.006 | 1.000 |
| BG-NM |  | -0.010 | 0.005 | 0.375 | -0.005 | 0.006 | 0.989 |
| WG-NM |  | 0.001 | 0.005 | 1.000 | 0.000 | 0.006 | 1.000 |
| CR-NM |  | -0.006 | 0.005 | 0.881 | -0.006 | 0.006 | 0.939 |
| GD-NM |  | -0.002 | 0.005 | 1.000 | 0.008 | 0.006 | 0.767 |
| AS-NM |  | -0.008 | 0.005 | 0.607 | -0.007 | 0.006 | 0.838 |
| PG-NM | sunny | -0.009 | 0.005 | 0.612 | -0.012 | 0.006 | 0.288 |
| AL-NM |  | -0.019 | 0.005 | **0.002** | -0.022 | 0.006 | **0.002** |
| BL-NM |  | -0.003 | 0.005 | 0.998 | -0.007 | 0.006 | 0.925 |
| WL-NM |  | -0.013 | 0.005 | 0.103 | -0.015 | 0.006 | 0.062 |
| BG-NM |  | -0.003 | 0.005 | 0.999 | -0.007 | 0.006 | 0.886 |
| WG-NM |  | 0.006 | 0.005 | 0.957 | -0.001 | 0.006 | 1.000 |
| CR-NM |  | -0.019 | 0.005 | **0.003** | -0.020 | 0.006 | **0.006** |
| GD-NM |  | 0.000 | 0.005 | 1.000 | 0.007 | 0.006 | 0.891 |
| AS-NM |  | 0.005 | 0.005 | 0.960 | 0.000 | 0.006 | 1.000 |

Table S3

| **Background matching effect of colour (dS) pairwise comparison between the pattern treatments.** | | | | | | | |
| --- | --- | --- | --- | --- | --- | --- | --- |
| contrast | Vegetation | dog | | | cat | | |
|  |  | estimate | SE | p.value | estimate | SE | p.value |
| BE-NM | B | 0.456 | 0.123 | **0.004** | 0.308 | 0.073 | **0.001** |
| BG-NM |  | 0.204 | 0.124 | 0.814 | 0.141 | 0.074 | 0.598 |
| GD-NM |  | 0.348 | 0.128 | 0.101 | 0.318 | 0.076 | **0.001** |
| NM-PB |  | -0.265 | 0.125 | 0.422 | -0.104 | 0.074 | 0.941 |
| NM-PG |  | -0.224 | 0.150 | 0.900 | -0.202 | 0.089 | 0.319 |
| NM-WE |  | -0.454 | 0.124 | **0.004** | -0.347 | 0.074 | **<.0001** |
| NM-WG |  | -0.088 | 0.125 | 1.000 | -0.097 | 0.074 | 0.966 |
| AS-NM | F | -0.005 | 0.123 | 1.000 | 0.001 | 0.073 | 1.000 |
| BE-NM |  | 0.101 | 0.123 | 1.000 | 0.053 | 0.073 | 1.000 |
| BG-NM |  | 0.062 | 0.124 | 1.000 | 0.061 | 0.074 | 1.000 |
| GD-NM |  | -0.097 | 0.127 | 1.000 | 0.075 | 0.075 | 0.998 |
| NM-PB |  | -0.226 | 0.125 | 0.686 | -0.095 | 0.074 | 0.972 |
| NM-PG |  | 0.200 | 0.150 | 0.960 | 0.016 | 0.089 | 1.000 |
| NM-WE |  | 0.174 | 0.124 | 0.940 | 0.052 | 0.074 | 1.000 |
| NM-WG |  | 0.064 | 0.125 | 1.000 | -0.032 | 0.074 | 1.000 |

Table S4

| **Background matching effect of luminance channel (dS_Lum_) pairwise comparison between the pattern treatments and environmental factors.** | | | | | | |
| --- | --- | --- | --- | --- | --- | --- |
|  | dog | | | cat | | |
| contrast | estimate | SE | p.value | estimate | SE | p.value |
| BE-NM | 0.072 | 0.044 | 0.784 | 0.077 | 0.046 | 0.758 |
| BG-NM | 0.086 | 0.044 | 0.584 | 0.093 | 0.046 | 0.518 |
| PB-NM | **0.276** | 0.045 | **<0.001** | **0.287** | 0.046 | **<0.001** |
| PG-NM | 0.036 | 0.057 | 0.999 | 0.038 | 0.059 | 0.999 |
| WE-NM | -0.030 | 0.044 | 0.999 | -0.030 | 0.046 | 0.999 |
| WG-NM | -0.034 | 0.045 | 0.998 | -0.034 | 0.046 | 0.998 |
| GD-NM | **-0.175** | 0.046 | **0.005** | **-0.181** | 0.047 | **0.005** |
| AS-NM | 0.110 | 0.044 | 0.239 | 0.115 | 0.046 | 0.224 |
| cloudy-sunny | -0.020 | 0.022 | 0.386 | -0.019 | 0.023 | 0.416 |
| B-F | **-0.059** | 0.022 | **0.006** | **-0.061** | 0.022 | **0.006** |
